# Deep clustering of protein folding simulations

**DOI:** 10.1101/339879

**Authors:** Debsindhu Bhowmik, Shang Gao, Michael T Young, Arvind Ramanathan

**Author notes:** Correspondence Computational Science and Engineering Division, Oak Ridge National Laboratory, One Bethel Valley Road, MS6085, Oak Ridge, Tennessee, United States Full list of author information is available at the end of the article.

## Abstract

We examine the problem of clustering biomolecular simulations using deep learning techniques. Since biomolecular simulation datasets are inherently high dimensional, it is often necessary to build low dimensional representations that can be used to extract quantitative insights into the atomistic mechanisms that underlie complex biological processes. In this paper, we use a convolutional variational autoencoder (CVAE) to learn low dimensional, biophysically relevant latent features from long time-scale protein folding simulations in an unsupervised manner. We demonstrate our approach on three model protein folding systems, namely the Fs-peptide (14 *μ*s aggregate sampling), villin head piece (single trajectory of 125 *μ*s) and the mixed **β*-*β*-*α** (BBA) protein (223 + 102 *μ*s sampling across two independent trajectories). In these systems, we show that the CVAE latent features learned correspond to distinct conformational substates along the protein folding pathways. The CVAE model predicts nearly 89% of all contacts within the folding trajectories correctly, while being able to extract folded, unfolded and potentially misfolded states in an unsupervised manner. Further, the CVAE model can be used to learn latent features of protein folding that can be applied to other independent trajectories, making it particularly attractive for identifying intrinsic features that correspond to conformational substates that share similar structural features. Together, we show that the CVAE model can quantitatively describe complex biophysical processes such as protein folding.

## 1 Introduction

The phenomenal growth of computing capabilities have accelerated our ability to precisely model and understand complex bio-molecular events at the atomistic scale [1, 2, 3]. Several recent studies have demonstrated how long timescale molecular dynamics (MD) simulations can provide detailed insights into events driving several complex biological phenomena such as protein folding, ligand binding, and membrane transport, often complementing experimental results. MD simulations are governed by a potential energy function that includes both bonded and non-bonded terms whose gradient defines a force-field applied to every atom in the biomolecular system [4]. These simulations integrate Newton’s laws of motion for every atom in the system using time-steps that typically are of the order of a femtosecond (10^−15^s). Even small simulation systems can potentially consist of thousands of atoms; given that bio-molecular events of interest typically occur at micro- and milli-second timescales, the increase in the size and complexity of these simulations is quickly becoming a limiting factor for extracting quantitative insights that are also biologically meaningful [5].

To overcome this challenge, a number of machine learning (ML) techniques are being developed to extract quantitative, biophysically relevant information from MD simulations. In particular, machine learning tools are able to quantify statistical insights into the time-dependent structural changes a biomolecule undergoes in simulations, identify events that characterize large-scale conformational changes at multiple timescales, build low-dimensional representations of simulation data capturing biophysical/biochemical/biological information, use these low-dimensional representations to infer kinetically and energetically coherent conformational sub-states, and obtain quantitative comparisons with experiments [6].

Since the dimensionality of MD simulations is large (3 × *N*, where *N* is the number of atoms, or 2 × (*φ,ψ,χ*) dihedral angles in the system of interest), ML techniques have focused on building low-dimensional representations of MD simulations. These dimensionality reduction techniques have used linear (e.g., principal component analysis [7], anharmonic conformational analysis [8, 9, 10]), non-linear (e.g., isometric mapping/ isomap [11], diffusion maps [12]) or hybrid approaches (e.g., locally linear embedding [13]) to characterize the conformational landscape sampled within simulations. Traditional ML approaches for analyzing long time-scale simulations typically require well-designed and often hand-crafted features. This in turn requires extensive prior knowledge about the system that is being simulated (for e.g., biophysically relevant reaction coordinates such as contacts between a ligand and its receptor). Often, use of certain ML techniques artificially restrict the simulation data being examined (for e.g., isolating only a subset of atoms from the simulations), or be prohibitively expensive to pre-/post-process the data. Finally, many of these approaches also require pairwise comparison of individual conformers within the simulation, with a similarity/dissimilarity measure that may be computationally expensive.

Deep structured learning approaches, on the other hand, overcome these challenges by automatically learning lower-level representations (or features) from the input data and successively aggregating them such that they can be used in a variety of supervised, semisupervised and unsupervised machine learning tasks [14]. Deep learning techniques have proven useful for a variety of structural bioinformatics applications, including protein structure prediction from biological sequences, and virtual screening/drug discovery applications [15, 16, 17]. Doerr and colleagues evaluated a variety of dimensionality reduction techniques for MD simulations and demonstrated that a shallow auto encoder could be used to visualize folding events within protein folding trajectories [18]. More recently, Pande and colleagues demonstrated how a reduced dimensionality representation from simulations built using tICA could be propagated using a time-dependent variational auto-encoder [19].

In this paper, we develop a convolutional variational auto-encoder (CVAE) that can automatically reduce the high dimensionality of protein folding trajectories and automatically cluster conformations from MD simulations into a small number of conformational states that share similar structural, and energetic characteristics. Using equilibrium folding simulations of Fs peptide, villin headpiece, and BBA, all model systems for protein folding, our CVAE discovers latent features that automatically captures folding intermediate states, including misfolded states that can be challenging to characterize. We further demonstrate that the learned latent features from the CVAE can be ‘transferred’ across simulations, making it relevant for succinctly summarizing large-scale simulations and compare behaviors across trajectories. Together, we show that deep learning techniques can be used for unsupervised learning of biophysically relevant latent features from long timescale MD simulations.

## 2 Methods

### 2.1 Datasets and Pre-processing

Given that deep learning techniques require large training data, we used three available datasets to demonstrate our approach. The first dataset consists of 28 separate MD simulations of the Fs peptide (Ace-A_5_(AAARA)_3_A-NME; 21 residues), a model system for protein folding, resulting in an aggregate sampling of 14 μs, consisting of 280,000 conformations. All simulations were performed at 300K using implicit solvent GBSA-OBC potentials and the AMBER-FF99SB-ILDN force field. This dataset was obtained from the MSMBuilder2 software [20]. The second dataset consists of (i) a single MD run of the Nle/Nle double variant of the C-terminal fragment of the villin head piece (referred to as VHP in this paper; PDB ID 2F4K) of 125 *μ*s simulated at 360 K, and (ii) two long MD runs of the mixed **β*-*β*-*α** fold, namely BBA (PDB ID: 1FME; 28 residues) for about 223 μs and 102μs at 325 K using Anton, a special purpose supercomputer for MD simulations and the CHARMM22* force field and a modified TIP3P water model compatible with CHARMM force field. For the VHP system, For the BBA system, we used a total of 1.1 million conformations of which 0.99 million conformers were used for training, with 0.01 million conformers used for testing/validation [21]. We used the second trajectory from the BBA simulations for testing our model based on the training from the first long time-scale trajectory.

We processed each trajectory using the `MDAnalysis` library [22, 23] to extract contact matrices between every pair of C^**α**^ atoms; we consider an atom to be in contact to another atom if it is separated by less than an 8 Å. Note that contact matrix representation is independent of rotation/translation (which is typically an artifact of MD simulations).

### 2.2 Convolutional Variational Autoencoder (CVAE)

Autoencoders (AEs) are a deep learning architecture designed to capture key representational information for a dataset within a low-dimensional latent space in an unsupervised fashion [24]. Autoencoders typically have an hourglass shaped architecture in which data is compressed into a low-dimensional latent space in the early layers and then reconstructed back in later layers. Therefore, the latent space learns to capture the most essential information required for reconstruction.

In variational autoencoders (VAEs), an additional optimization constraint is added that requires the latent space to be normally distributed [25]. While regular autoencoders may effectively capture important information in a reduced dimension, the latent embeddings may be sparsely distributed; this typically means that key information is spread across several clusters in the latent space, and the empty space between clusters does not capture any useful information-sampling from this empty space typically creates nonsensical results. By forcing the latent space to be normally distributed, we force the network to fully utilize the latent space so that information is distributed more evenly; this allows us to sample from any point in the latent space to generate new results that reflect the patterns in the original dataset. We therefore chose to use a VAE instead of a regular AE, as one of our long-term goals is to generate new potential structures based on the information learned within our model.

Rather than using regular feedforward layers in our VAE, we apply convolutional layers because they utilize sliding filter maps that can better recognize local patterns independent of its position in the data. In contact map representations, the state of the protein depends on the local interactions between a few atoms rather than on the global position of all atoms in the protein. Because these local interactions do not always appear in the exact same place in the protein, convolutional layers are better suited to recognize these local patterns independent of their position compared to feedforward networks. The architecture for the convolutional autoencoder (CVAE) used in our experiments is illustrated in Fig. 1.

**Figure 1.**
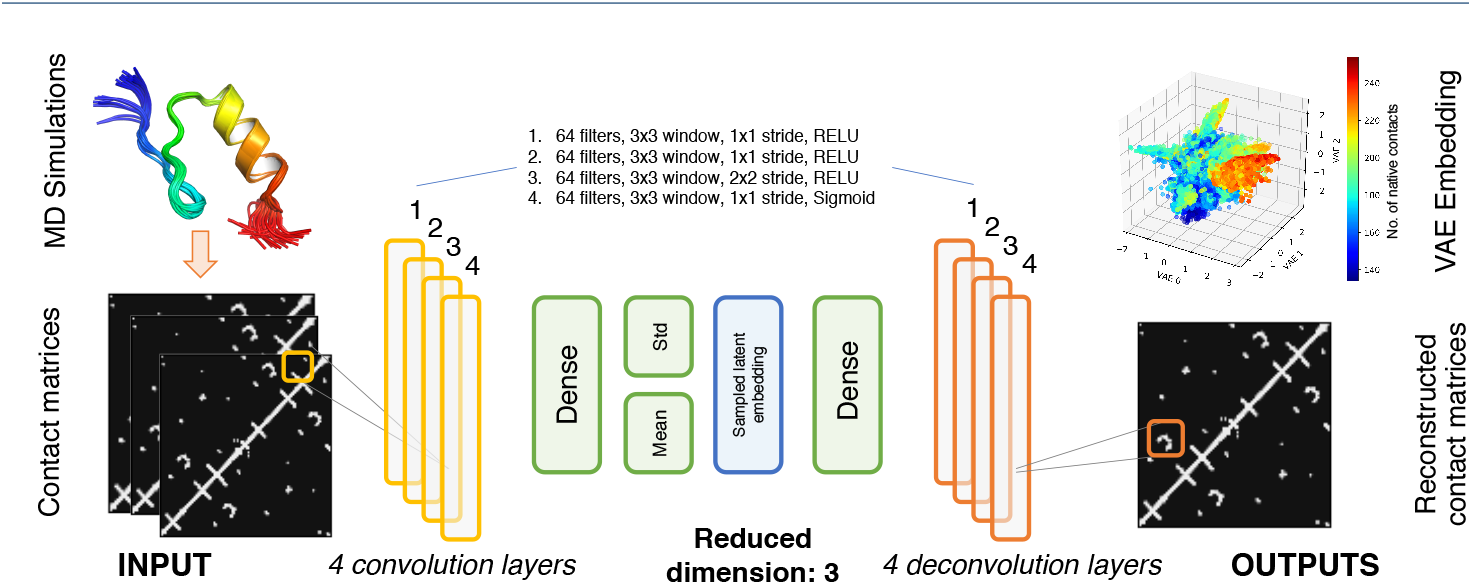
Convolutional variational autoencoder architecture. The deep learning network processes MD simulation data into contact maps (2D images) that are then successively fed into 4 convolutional layers. The outputs from the final convolutional layer is then fed into a fully connected (dense) layer. This is then used to build the latent space in three dimensions, the output of which is the learned VAE embedding. In order to reconstruct the contact maps, we then use 4 successive de-convolutional layers, symmetric to the 4 input convolutional layers.

Each CVAE was trained for a fixed number of epochs that was determined by the convergence of loss and variance-bias trade-off. The batch size was selected to be relatively small (length of the training data/100) to ensure reduced data in latent space do not collapse. We divided each dataset into training/testing (80%/20% of the simulation trajectories) – although not a requirement for unsupervised learning techniques, we used the testing data to characterize both the clustering and reconstruction quality of the CVAE. The various hyperparameter settings are shown in Table 1

**Table 1.** Hyperparameter settings used for CVAE training

| Hyperparameter | Lower Bound | Upper Bound | Optimal Fs-Peptide | Optimal BBA | Optimal VHP |
| --- | --- | --- | --- | --- | --- |
| Num Convolutional Layers | 1 | 4 | 4 | 4 | 4 |
| Num Convolutional Filters | 16 | 125 | 100 | 100 | 99 |
| Convolutional Filter Shape ($n \times n$) | 2 | 7 | 5 | 5 | 5 |
| Num Dense Neurons | 32 | 100 | 64 | 64 | 58 |
| Latent Dimension | 2 | 16 | 10 | 10 | 9 |
| Mean Reconstruction Error |  |  | 7.82 | 24.31 | 74.66 |
| Mean Reconstruction Error (latent dim 3) |  |  | 23.24 | 53.48 | 113.07 |

## 3 Results

We posited that the CVAE (described in Section 2) encoding would result in a model that can automatically capture biophysically relevant features from the simulation datasets. We used three model protein folding systems, namely Fs peptide, villin headpiece (VHP) and BBA to demonstrate that the CVAE can learn a biophysically relevant latent space that corresponds to folding reaction coordinates, including fraction of native contacts and root mean squared deviation (RMSD) to the native state. To calculate the fraction of native contacts we use a definition similar to Savol and Chennubhotla [26]. Native contacts are based on a distance cut off of 8 A or less between between C^**α**^ atoms and at least 75% of conformations remain within an RMSD cut-off of 1.1 A of the native structure. First, we evaluate the ability of CVAE to learn a reduced dimensional space given the MD simulation data. Second, we show that the CVAE latent space corresponds to biophysically relevant features for each of the folding simulations studied. Finally, we demonstrate that the CVAE latent features can be transferred to other simulations, making it generalizable to a particular protein type.

### 3.1 Reconstruction quality of CVAE on protein folding trajectories

In order to evaluate the CVAE reconstruction quality from the protein folding trajectories, we first examined the overall loss 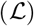 of the CVAE over the training epochs (Equation 1).

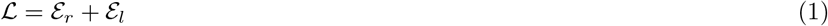

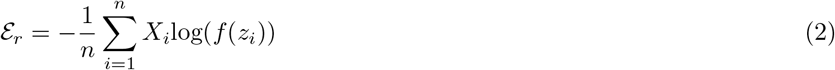

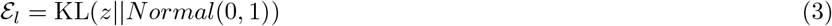

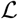 is characterized as the sum of the reconstruction loss 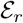 (Equation 2) and the latent loss 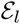 (Equation 3). The reconstruction loss measures how well the CVAE can reconstruct the original input and is calculated as the cross entropy loss between *f*(*z*), which indicates the reconstructed probability of contact between two C^**α**^ atoms, and the original *X* conformations from the simulation, which indicate the existence of contact between two C^**α**^ atoms. The latent loss is a regularizing constraint that forces the latent embeddings *z* to conform to a Gaussian distribution; this is calculated as the Kullback-Leibler (KL) divergence between the latent embeddings *z* and a Normal distribution with mean 0 and standard deviation 1.

For the three protein folding trajectories in this study (Figure 2)A-C, we observed that the overall loss, 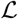, stabilizes over the training epochs, showing that it converges over time. We observed that for each protein, the number of training epochs needed to reach convergence is different; this is not surprising, given that the size of these proteins are different and the trajectories have unique folding pathways. Furthermore, we observed that the reconstruction loss (described in Equation 1) is also different for each protein system – indicating that the quality of CVAE reconstruction is unique to each protein system.

**Figure 2.**
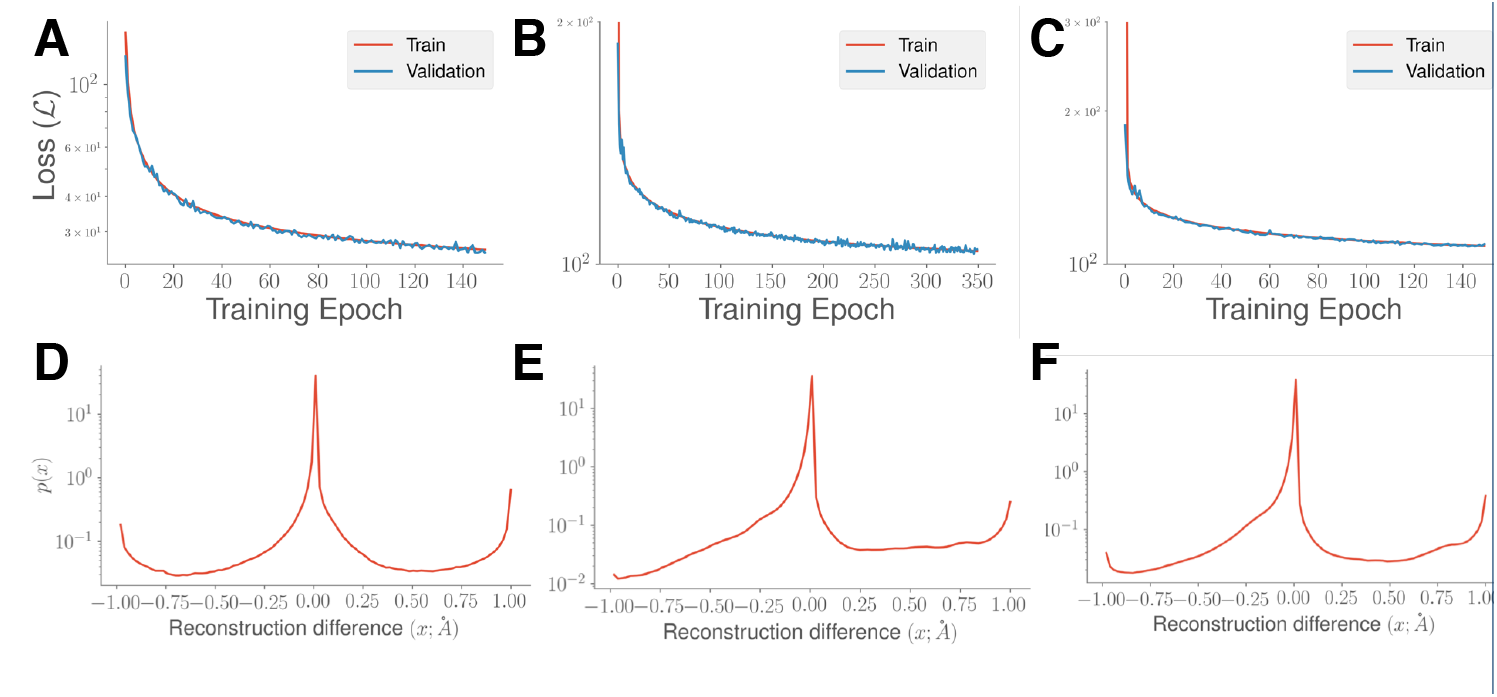
Quantifying CVAE performance over MD simulations. We quantify the loss, 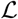, described in the Results section over the number of training epochs for each of the systems, namely (A) Fs-peptide, (B) VHP, and (C) BBA. As the number of epochs increases, the loss function decreases, indicating convergence in the training process. We also quantify the quality of reconstruction based on the CVAE’s ability to predict contacts within the testing dataset. For each of the simulations, namely (D) Fs-peptide, (E) VHP, and (F) BBA, we show the reconstruction difference between the actual and predicted contact matrices from the testing data and histogram it to quantify how many contacts may be mispredicted by the model. See text for additional details.

We next examined whether the CVAE latent features can faithfully reconstruct the original data. To evaluate this, we used the reconstruction difference that simply measures the difference between the reconstructed and original contact matrices. Note that while the original contact matrix is typically binary (indicating presence or absence of a contact), the output from the CVAE is a value between 0 and 1, which may be interpreted as a likelihood of observing a contact between two C^**α**^ atoms. We plotted a histogram of reconstruction from the testing data, shown in Figure 2D. The reconstruction difference varies between −1 and +1, which indicates whether the reconstructed data mispredicts the presence or absence of a contact respectively. We choose a nominal threshold of 10% of the original value to indicate misprediction. For the Fs-peptide simulations, the CVAE is able to faithfully reconstruct nearly 88% of all the observed contacts and mispredicts only 12% of contacts (Figure 2D). We note that these contacts are at the interfaces of secondary structural elements, between *α*-helices, or between *α*-helices and *β*-strands. We can make similar observations for the other protein systems; the average reconstruction error for VHP is about 10.6% (Figure 2E) and for BBA (Figure 2F), the CVAE reconstruction can recover nearly 88.5% of all contacts correctly in the folding simulations.

We evaluated the performance of CVAE as a function of several model hyperparameters using Bayesian optimization [27, 28, 29]. The search bounds and optimal results for the hyperparameters are summarized in Table 1. While the optimal settings for the latent dimension for each molecule was found to be near ten, we chose to use models with latent dimensions of size three. Since it is possible to verify visually that the autoencoder is meaningfully capturing the folding process without sacrificing much in terms of reconstruction error, it is reasonable to maintain a three dimensional latent space. The mean reconstruction loss over various settings for the latent dimension for each of the molecules can be seen in Figure 3. When considering the latent dimension, there exists a trade-off between the model’s ability to compress information and its ability to minimize reconstruction error. Further, we find that the model’s performance can be affected by the interactions between the choice of latent dimension and other model hyperparameters. We therefore found visualization techniques such as t-distributed stochastic neighborhood embedding (t-SNE) [30] valuable when verifying the choice for the size of the model’s latent dimension.

**Figure 3.**
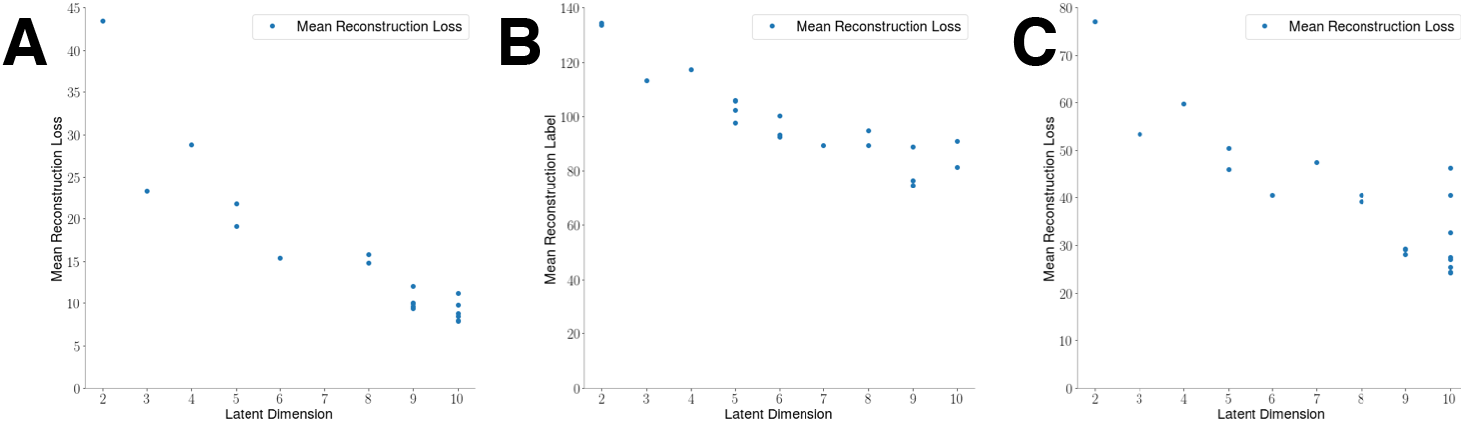
Mean reconstruction error over various latent dimensions indicates as the latent dimension increases, the mean reconstruction error decreases. This shows a trade-off between compression and model accuracy. (A) Fs-Peptide exhibits a linear decrease in reconstruction error as the latent dimension increases. (B) VHP begins to show the limits of increasing the model’s latent dimension; the lowest recorded mean reconstruction loss was found to be at latent dimension nine. (C) While the lowest reconstruction error for BBA was found to be with a latent dimension of size ten, there is a wide amount of variation in the reconstruction loss for this latent dimension size. Various values of the reconstruction error at each latent dimension shows there to be interactions between the various model hyperparameters.

### 3.2 CVAE reveals folding intermediates of Fs-peptide

Fs-peptide is often used as a model system to study protein folding processes. We examined whether our CVAE can recapitulate the diverse *α*-helical conformational substates in an unsupervised manner. Figure 4A shows the latent space learned by the CVAE. Each conformation from the training data is represented as a three-dimensional coordinate in the latent space. To understand whether the latent space captured by the CVAE describes the folding process, we colored each conformer with the corresponding RMSD to the native state. The RMSD to the native state is often used as a conformational coordinate to track protein folding trajectories [31]. We note that the input to the CVAE is only the raw contact maps; however, the model is able to distinguish between low and high RMSD conformers when projected onto the latent space.

**Figure 4.**
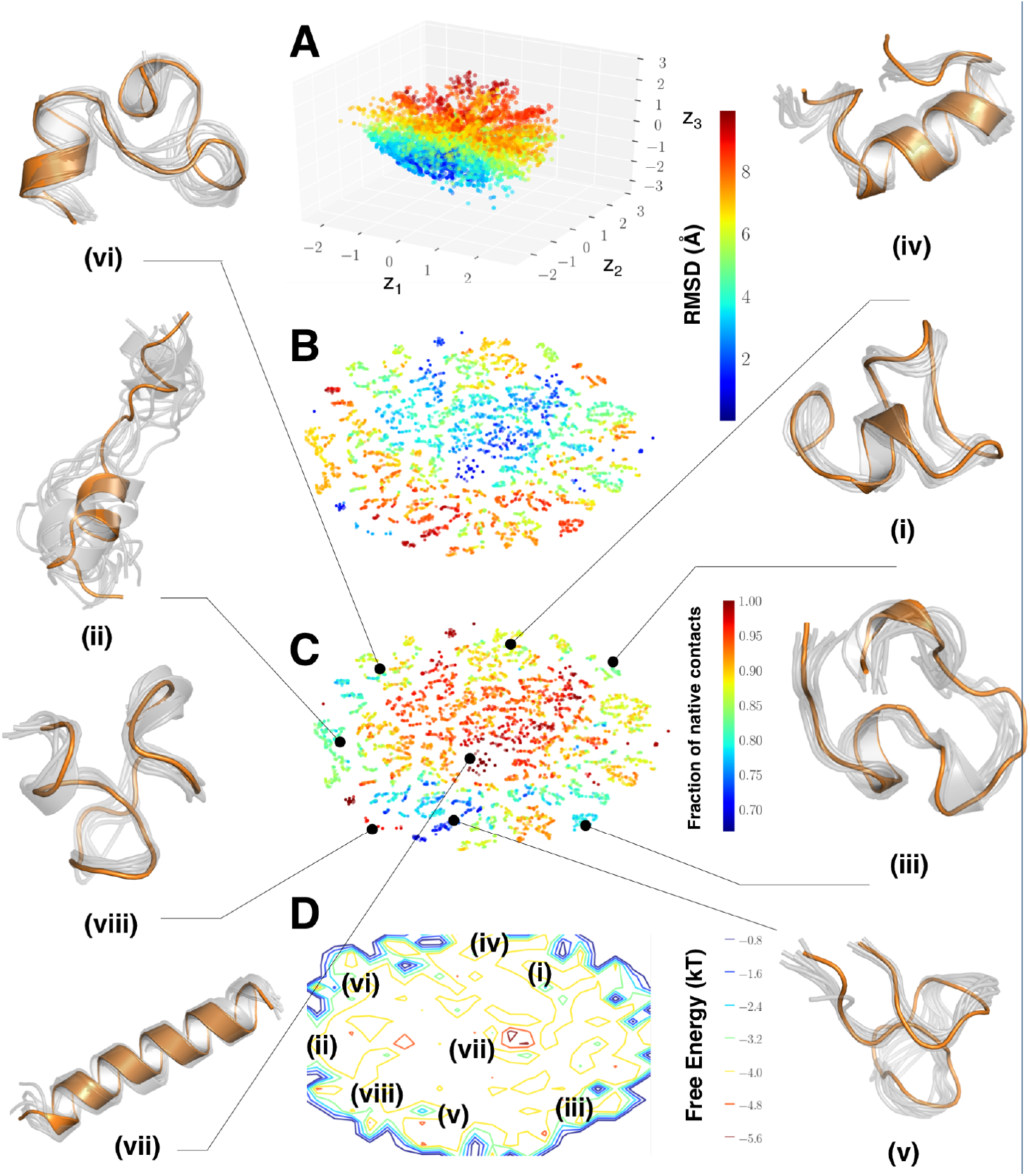
CVAE can identify conformational substates from Fs-peptide simulations. (A) We project the conformations from the simulations onto a latent space spanned by the three dimensions of the CVAE (*z*_1_ – *z*_3_). Each conformation is then painted with the corresponding RMSD to the native state. As observed from the figure, there is a gradation from unfolded states (i.e., higher RMSD values) towards more folded states (lower RMSD values). To allow easy interpretation, we use t-SNE to project the conformations. (B) t-SNE projection painted with RMSD values and (C) fraction of native contacts shows clear separation between folded conformations, labeled (vii) along with partially folded states, namely (ii), (iii), (iv), and (vi), along with unfolded states, labeled (v). It is notable that state (viii) has lower RMSD to the folded state (vii), where as shows very few of the native contacts that are found in the folded state, implying that this state may be a misfolded state in the ensemble simulations. (D) A simple histogram representation of the projected t-SNE space allows us to characterize the conformational states observed within the simulations. The labels provide a relative description of the location of the various conformational states.

Within the latent space, we note the presence of distinct pockets with high RMSD values to the native state (red colors), which converge eventually into folded state (blue colors). The gradation of the colors along the arms of the CVAE axes indicates that the latent space (*z*_1_ – *z*_3_) is able to describe the folding process. It indicates multiple pathways along which Fs-peptide folds into its final state. Although the CVAE-determined latent space can capture the presence of both folded/unfolded states (quantified by the total number of contacts), it is still challenging to interpret. Hence, we used t-SNE to visualize the results. We painted each conformation in the t-SNE with the RMSD values (Figure 4B) and the fraction of native contacts (Figure 4C). Even for a The t-SNE approach allows us to separately identify distinct conformational observed from the simulations, labeled (i) to (viii), in the folding trajectories. In particular, we find the presence of partially folded *α*-helical bundles as well as a fully formed *α*-helix, which represents the folded state of the protein. Additionally, we also find that the different folded states are separated and connected via multiple intermediate states, all of which have relatively lower number of total contacts. This indicates that for the transitions between the folded microstates, the peptide must undergo several unfolding events.

Interestingly, our approach also reveals the presence of potentially misfolded states in these trajectories. In this work, we consider a misfolded state to be a set of conformations that share higher fraction of native contacts, but have a high RMSD from the native state ensemble of the protein. For example, state (viii) in Figure 4C shows the presence of conformations that have higher fraction of native contacts (close to 0.95), however, its secondary structure content is significantly different from the native state, highlighted as (vii) in Figure 4C. The intermediate states identified here have differences in their secondary structural content, i.e., the number of *α*-helical turns as depicted in states labeled (i), (ii), (iv), (vi) and (viii) along with differences in the extent to which the N- and C-terminal ends of the protein are folded (for e.g., state labeled (ii) folds from the N-terminal end versus state labeled (iii) folds from the C-terminal end).

We can also visualize the tSNE dimensions as the logarithm of the histograms as a simple estimate of the free energy surface as depicted in Figure 4D, where by conformational states can be visualized. This representation is only for visual purposes and as such can be used for qualitative insights into the organization of the folding energy landscape of Fs-peptide. The native state of the protein, labeled (vii), consists of the fully folded peptide, while many of the partially folded states and their intermediates are distributed around the periphery of this landscape. It is interesting to note that the contours represent conformational states that correspond to folding coordinates and each of the states are marked (using solid lines) as to where they belong on this landscape.

### 3.3 CVAE reveals conformational states in the VHP folding pathway

For the VHP simulations, we were able to identify a similar distribution of folded/unfolded conformations along its folding pathway (Figure 5). Even though the reconstruction error plots from Figure 3B indicate that the ideal number of latent dimensions is 9, we examined whether a low dimensional encoding with just three dimensions is able to capture folding events within this trajectory. Similar to the analysis of the Fs-peptide folding trajectories, the latent embedding of the CVAE reveals the presence of folded and unfolded conformations that are separated by a a large number of intermediate states (Figure 5A). Since these simulations were carried out at a higher temperature (360 K), these simulations indicate larger fluctuations in the secondary structures of VHP. Further, within the course of the simulations, a total of 34 folding events are summarized, which indicate a large number of conformational states actually correspond to folded conformations.

**Figure 5.**
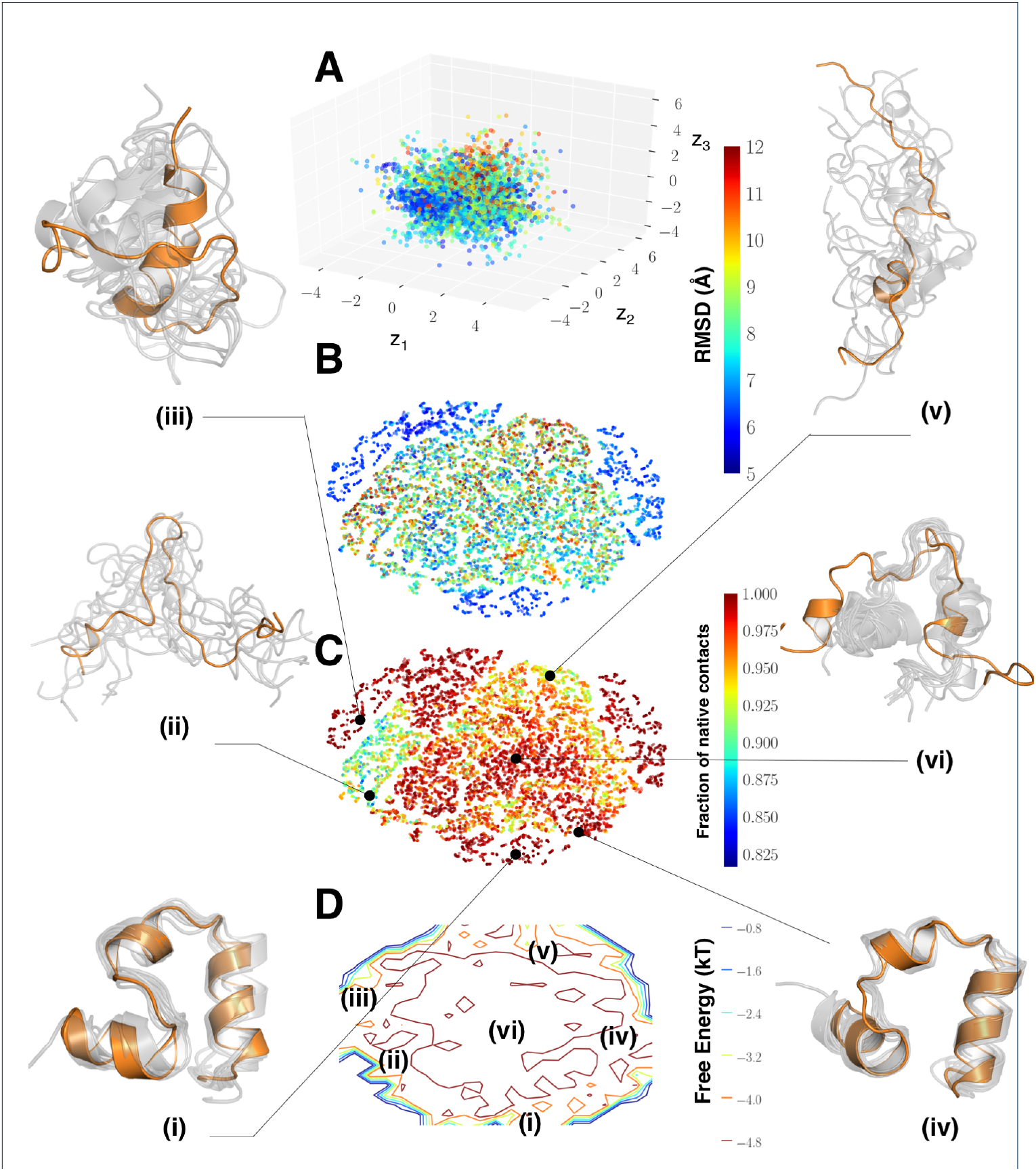
CVAE identifies partially folded states within the VHP simulations. (A) For the VHP simulations, we project the conformations onto the CVAE space spanned by the three latent dimensions (*z*_1_ – *z*_3_). Each conformation is painted by the RMSD to the native state with the color scale depicting the range of RMSD values between the folded and unfolded states. For the VHP simulations we consider any state with an RMSD of less than 5 A as essentially folded (since these simulations are carried out at a higher temperature). To enable a better characterization of the VHP folding landscape, we project the conformations using t-SNE to identify conformational substates. Each conformation is painted with (B) the RMSD to the native state, and (C) the fraction of native contacts. The separation between the folded, unfolded and intermediate states can be understood by examining the conformational states labeled (i) - (v). (D) A simple histogram representation from the projected t-SNE space allows us to characterize the locations of the conformational states observed from this simulation. The labels provide a relative location of the various conformational states.

To enable interpretation of the VHP folding landscape, we projected the CVAE latent dimensions using t-SNE and observed that the folded states of VHP are separated into three distinct ‘wells’ that correspond to the folding events along this trajectory. The evidence for the folding events emerges from painting the t-SNE landscape with the fraction of native contacts (Figure 5C). A large portion of the trajectory is either unfolded (e.g., states labeled (ii), (v) indicated as conformational ensembles along the trajectory) or partially folded, i.e., showing the presence of all the three helices, but with different number of helical turns (e.g., states labeled (iii) and (vi) in Figure 5). Finally, the folded states labelled (i) and (iv) capture distinct orientations of *α*-helices as observed from the figure. It is interesting to note that the transition from one folded state to the other involves partial unfolding (similar to Fs-peptide). Further, we also note that the partially folded state (iii) consists of many native contacts; however, this state does not have all the three helices and may represent an unfolded intermediate state through which the transition to either of the two folded states may occur. The simple histogram representation of the t-SNE coordinates (Figure 5D) provides an easy way to interpret the different conformational states with respect to the folded states in the trajectory (states (i) and (iv)).

### 3.4 CVAE analysis of BBA folding simulations can be transferred to learn folding patterns across trajectories

We next examined whether the CVAE learned features could be used to predict conformational states from a completely different trajectory. To facilitate this analysis, we used the BBA simulations (see Methods section) as a prototype example. Our experimental set up included training the CVAE on the first long time-scale trajectory (223 *μ*s) and predicting if it captures the folding events from the second trajectory (102 *μ*s). As depicted in Figure 6A, the three latent CVAE dimensions capture the presence of multiple folded conformational states (labeled (i), (iii) and (iv) in Figure 6B using t-SNE). These states are separated by an intermediate state labeled (ii) and an unfolded state labeled (vi). Finally, it is interesting to note that latent space also characterizes a misfolded state, labeled (v), which shows the presence of an extended **β**-strand.

**Figure 6.**
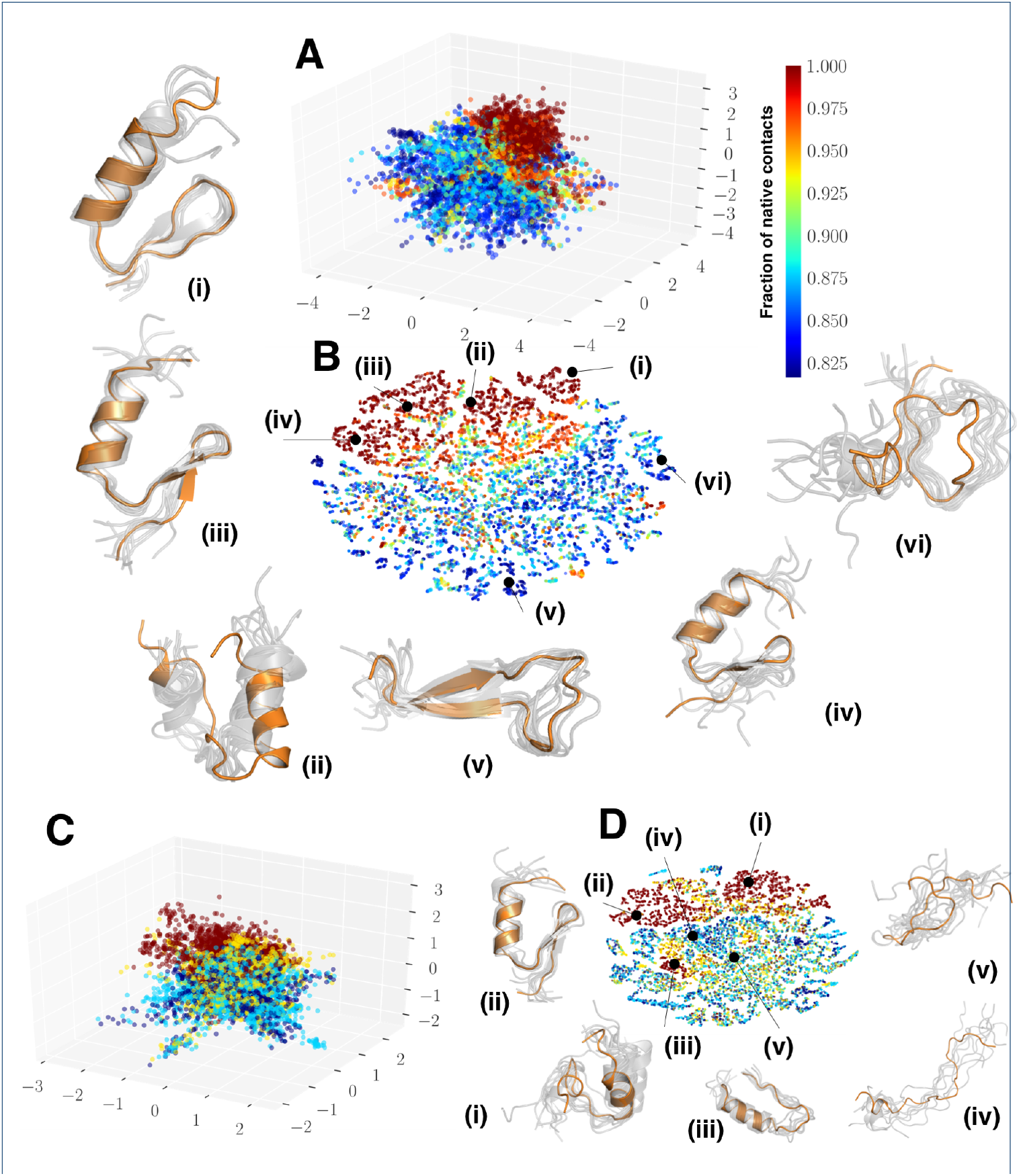
CVAE learned features predict folding intermediates across two independent folding simulations of BBA protein. (A) depicts the CVAE latent space embedding on one 223 *μ*s trajectory of BBA folding. Each conformation from the simulation is projected onto a three dimensional embedding and painted with the fraction of native contacts. One can observe that folded and unfolded states are separated into distinct clusters. (B) To elucidate the embedding from the CVAE, we project the conformations from the trajectory using t-SNE and identify conformational states captured by the CVAE. States are captured as described in Figure 4 and shown as a cartoon representation based on the cluster from which they belong to. These conformational states depict various levels of BBA folding, labeled (i) through (vi). (C) Using the model learned on the longer trajectory, we project the conformers from the second independent simulation of length 102 μs onto the same latent space. It is notable that the folded states are similarly clustered together while the unfolded states are captured separately. (D) The latent representation from the CVAE can be applied across different trajectories to summarize folded states. As shown in the cartoon representation, the conformational ensembles are separated into distinct folded (labeled (ii)), partially folded (labeled (i) and (iii)) and fully unfolded states (labeled (iv) and (v)).

Using the same model that was trained on the first trajectory, we can project the conformations from the second, shorter simulation onto the latent space learned to test if the folded/unfolded states are separated. As shown in Figures 6C and 6D, the latent space from the second trajectory clearly shows a separation between the folded states labeled (ii) in Figure 6D, partially folded states labeled (i) and (iii) in Figure 6D, and unfolded states labeled (iv) and (v) in Figure 6D. We also observed that the latent space reconstruction difference is on par with the original model, implying that the features learned by the CVAE can be transferred

## 4 Discussion/Conclusion

We have demonstrated how deep learning algorithms can be used to analyze and interpret protein folding simulations. We designed a CVAE that can encode the inherent high dimensionality of the folding trajectories into a low dimensional embedding that is biophysically relevant. We demonstrated our approach on three prototypical systems, namely Fs-peptide, VHP and BBA, all of which have been extensively characterized in previous studies. In all the cases, we note that the learned CVAE embeddings captured the distinction between potentially folded, partially folded, and misfolded states.

In this paper, we used contact matrices determined form the simulations as inputs to the CVAE. Contact matrices are a practical approach to represent simulation datasets, which have been widely used to characterize protein folding pathways [32, 33]. However, the resolution of information captured using contact maps is fairly low and may not be specific (e.g., distinguish between native, non-native and misfolded states). Although the CVAE identified the presence of folded/unfolded and misfolded states in the simulations, there is significant scope for directly using coordinate information (or other physical quantities such as dihedral angles) from simulations for characterizing these pathways.

Complementary to the approaches taken by Doerr and colleagues [18], we build an autoencoder; however augmenting it with a variational formulation allows us to obtain interpretable features from the latent space. As demonstrated in the three systems, the CVAE latent spaces capture a succinct model of protein folding with the ability to distinguish conformational substates that share similar structural features. We have yet to evaluate whether these substates share similar energetic profiles. Further, our CVAE can be used to potentially augment propagators in time [19] such that temporal correlations are captured within these trajectories.

The selection of the hyperparameters, such as the size/stride of convolutional filters and the dimensions of the latent space to embed the simulations were based on empirical evaluations. Ideally, the choice of the latent space representation should be a parameter that can be learned from the simulation data itself (instead of being specified by the user). Further, these latent dimensions should correspond to directions in the landscape that enable the bio-molecular system to sample folded/misfolded states, which has been previously demonstrated by pursuing higher order statistical dependencies in atomic fluctuations in the simulations [8, 5]. We plan to extend our CVAE to automatically learn and infer the latent dimensional space.

Further studies are essential in associating the biophysical relevance of the learned CVAE embeddings. Specifically, we have not evaluated whether the CVAE embeddings for these folding trajectories correspond to biophysical reaction coordinates, i.e., whether the unique directions proposed by the CVAE can ‘fold’ a protein system. Temporal correlations are known to significantly influence bio-molecular events [34]. Although we trained our model to include temporal information (i.e., frames for the training was based on successive conformations in the trajectory), the embeddings learned do not necessarily correspond to detectable bio-molecular events. For e.g., in a protein folding trajectory, a typical event corresponds to ‘whether a *β*-strand was formed’ – our CVAE is currently unable to identify time-points where significant structural or dynamical changes have occurred within trajectories. Leveraging our previous experience in developing techniques for event detection [10, 35], we will explore deep learning models for bio-molecular event detection in the near future.

## Author’s contributions

AR and DB conceived and designed the study. DB and SG implemented the CVAE model. MTY contributed to developing tools for analyzing hyperparameters and their impact on CVAE performance. All authors contributed to the writing, editing and reviewing the article.

## Acknowledgements

The authors would like to thank D. E. Shaw Research for providing access to the molecular dynamics simulation trajectories of BPTI and fast protein folding simulations.

This manuscript has been authored by UT-Battelle, LLC under Contract No. DE-AC05-00OR22725 with the U.S. Department of Energy. The United States Government retains and the publisher, by accepting the article for publication, acknowledges that the United States Government retains a non-exclusive, paid-up, irrevocable, world-wide license to publish or reproduce the published form of the manuscript, or allow others to do so, for United States Government purposes. The Department of Energy will provide public access to these results of federally sponsored research in accordance with the DOE Public Access Plan (http://energy.gov/downloads/doe-public-access-plan).

This work has been supported in part by the Joint Design of Advanced Computing Solutions for Cancer (JDACS4C) program established by the U.S. Department of Energy (DOE) and the National Cancer Institute (NCI) of the National Institutes of Health. This work was performed under the auspices of the U.S. Department of Energy by Argonne National Laboratory under Contract DE-AC02-06-CH11357, Lawrence Livermore National Laboratory under Contract DE-AC52-07NA27344, Los Alamos National Laboratory under Contract DE-AC5206NA25396, Oak Ridge National Laboratory under Contract DE-AC05-00OR22725, and Frederick National Laboratory for Cancer Research under Contract HHSN261200800001E.

This research used resources of the Oak Ridge Leadership Computing Facility at the Oak Ridge National Laboratory, which is supported by the Office of Science of the U.S. Department of Energy under Contract No. DE-AC05-00OR22725.

